# The wheat dioxygenase BX6 is involved in the formation of benzoxazinoids *in planta* and contributes to plant defense against insect herbivores

**DOI:** 10.1101/2021.09.25.461767

**Authors:** Reut Shavit, Zhaniya S. Batyrshina, Beery Yaakov, Matilde Florean, Tobias G. Köllner, Vered Tzin

## Abstract

Benzoxazinoids are plant specialized metabolites with defense properties, highly abundant in wheat (*Triticum*), one of the world’s most important crops. The goal of our study was to characterize dioxygenase *BX6* genes in tetraploid and hexaploid wheat genotypes and to elucidate their effects on defense against herbivores. Phylogenetic analysis revealed four *BX6* genes in the hexaploid wheat *T. aestivum*, but only one ortholog was found in tetraploid (*T. turgidum*) wild emmer wheat and the cultivated durum wheat. Transcriptome sequencing of durum wheat plants damaged either by aphids or caterpillars revealed that several *BX* genes including *TtBX6* were upregulated upon caterpillar feeding relative to undamaged control plants. A virus-induced gene silencing approach was used to reduce the expression of *BX6* in *T. aestivum* plants and exhibited both reduced transcript levels and reduced accumulation of different benzoxazinoids. To elucidate the effect of *BX6* on plant defense, bioassays with different herbivores feeding on *BX6*-silenced leaves were conducted. The results showed that plants with silenced *BX6* were more susceptible to aphids and the two-spotted spider mite compared to controls. Overall, our study indicates that wheat BX6 is involved in the formation of benzoxazinoids *in planta* and contributes to plant resistance against insect herbivores.

## Introduction

Plants in nature are continuously challenged by diverse insect herbivores and, in response, evolved various defense mechanisms to reduce insect damage and to preserve their own fitness. Insect infestation is initially recognized by plants through herbivore-associated and damage-associated molecular patterns [1], followed by the activation of signaling pathways and metabolic responses that can remain local, limited to the infestation site, as well as spreading systematically [2,3]. Such metabolic modifications involve a complex network that influences the plant’s central and specialized metabolic activities [1]. The specialized metabolites often act as feeding deterrents, reducing damage from herbivores, and thus maintaining plant fitness. One class of these metabolites is the benzoxazinoids (BXDs), which are highly abundant in many species of the grass family, including important crops such as maize, wheat, rye, and wild barley species [4–6]. Moreover, BXDs have also been described to sporadically occur in few dicot species [6,7]. The protective effects of BXDs are mainly due to their antifeedant properties, caused by the inhibition of insect digestive proteases responsible for detoxification and pest salivation [8], and by the high levels of the compound DIMBOA (2,4-dihydroxy-7-methoxy-1,4-benzoxazin-3-one) that mediates callose formation [9,10].

The biosynthesis of BXDs has been investigated over the last two decades due to extensive research in maize [7,11,12]. It starts from indole-3-glycerol phosphate, an intermediate of the tryptophan biosynthetic pathway, produced by indole-3-glycerol synthase (*IGPS*), which has recently been suggested as the branchpoint between the formation of free indole, the aromatic amino acid tryptophan, and BXD biosynthesis [13] (Figure 1). Indole-3-glycerol phosphate is converted to indole through the action of the indole synthase BX1 (benzoxazinoneless 1), followed by four cytochrome P450 enzymes, BX2-5, which introduce four oxygen atoms and form the core benzoxazinoid structure 2,4-dihydroxy-1,4-benzoxazin-3-one (DIBOA). DIBOA can subsequently be glycosylated to DIBOA-Glc by the glucosyltransferases BX8 and BX9. Hydroxylation of DIBOA-Glc catalyzed by the dioxygenase BX6 and subsequent methylation of the added hydroxyl group by the *O*-methyltransferase (OMT) BX7 lead to DIMBOA-Glc, the major BXD in many maize lines [11]–[14]. Three OMTs that methylate DIMBOA-Glc into 2-hydroxy-4,7-dimethoxy-1,4-benzoxazin-3-one glucoside (HDMBOA-Glc) were recently identified. They are located on maize chromosome 1 (*BX10, BX11*, and *BX12*) and share high sequence similarities [17,18]. Likewise, two additional enzymes, the 2-oxoglutarate-dependent dioxygenase BX13 and the OMT BX14, were recently shown to be involved in converting DIMBOA-Glc into HDM_2_BOA-Glc in maize [19]. Glucosylated BXDs are stored in the vacuole, where they are protected against β-glucosidases, which are usually found in the plastids or associated with the cell wall [11]. Upon demand, BXDs are deglucosilated through the activity of β-glucosidases and subsequently transported into the extracellular matrix. The unstable aglycones are can be further converted into degradation products such as 6-methoxy-benzoxazolin-2-one (MBOA) or benzoxazolinone (BOA) [20].

**Figure 1.**
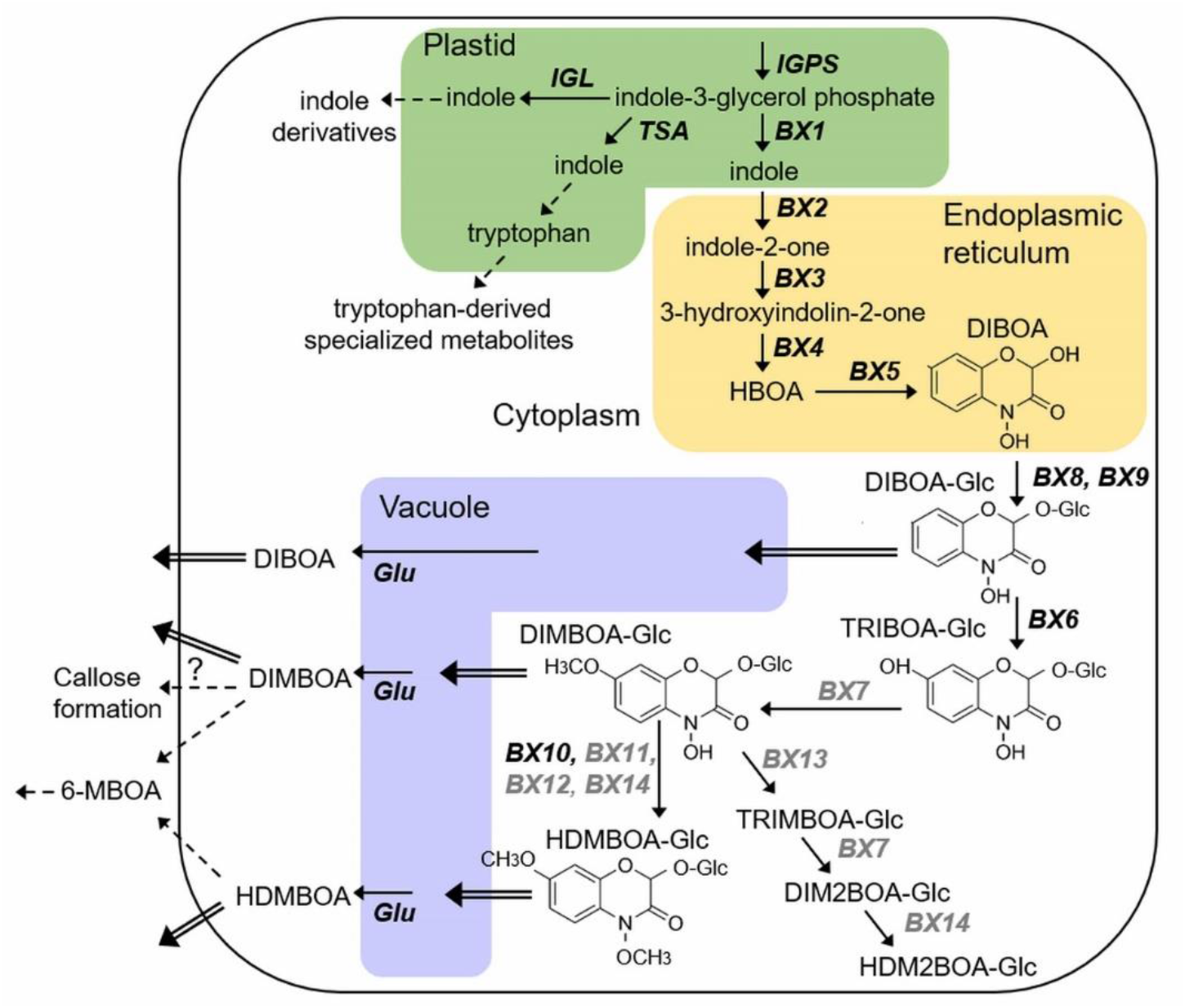
A schematic diagram of the indole and benzoxazinoid biosynthesis in maize. Genes that were characterized in both maize and wheat are written in black, while genes only identified in maize, to date, are written in grey. Dashed arrows represent multiple enzymatic steps. Double lines arrows represent the transportation of compounds into the vacuole for storage or to the extracellular matrix. Compound names: HBOA, 2-hydroxy-1,4-benzoxazin-3-one; DIBOA, 2,4-dihydroxy-1,4-benzoxazin-3-one; TRIBOA, 2,4,7-trihydroxy-2H-1,4-benzoxazin-3(4H)-one; DIMBOA, 2,4-dihydroxy-7-methoxy-1,4-benzoxazin-3-one; HDMBOA, 2-hydroxy-4,7-dimethoxy-1,4-benzoxazin-3-one; TRIMBOA, 2,4,7-trihydroxy-8-methoxy-1,4-benzoxazin-3-one; DIMB_2_BOA, 2,4-dihydroxy-7,8-dimethoxy-1,4-benzoxazin-3-one; HDM_2_BOA, 2-hydroxy-4,7,8-trimethoxy-1,4-benzoxazin-3-one; MBOA, 6-methoxy-benzoxazolin-2-one; Glc, glycosylated form of each molecule. Gene names: IGPS, indole-3-glycerolphosphate synthase, TSA, tryptophan synthase alpha subunit; IGL, indole-3-glycerolphosphate lyase; GLU, beta-glucosidase. Methylation of DIMBOA-Glc to HDMBOA-Glc in maize is catalyzed by three closely related OMTs, ZmBX10–ZmBX12, while in wheat, the appropriate enzyme is unrelated and named TaBX10.

Wheat, a staple crop that provides 20% of the world population’s caloric and protein intake, is well adapted to arid climates [21]. It accumulates large amounts of BXDs as defense compounds against herbivores and pathogens [16]. Orthologues of the maize genes *BX1-BX5* and *BX8-BX9*, which encode the core BXD pathway, have been identified in hexaploid bread wheat (BBAADD), tetraploid wheat (BBAA), and in the three diploid progenitors of hexaploid wheat (AA, BB, and DD subgenomes; [22,23]). Although DIMBOA-Glc and HDMBOA-Glc are abundant in wheat [24–26], orthologous genes of *BX6* and *BX7* have not been fully characterized. Very recently, Sue and coworkers described the discovery and enzymatic function of four BX6 isoforms in hexaploid *T. aestivum* [27]. However, the BX6 orthologs in tetraploid wheat and diploid wheat progenitors are still unknown. The recent identification of TaBX10, an OMT that methylates DIMBOA-Glc to HDMBOA-Glc in wheat, showed that TaBX10 and maize BX10-12 are not related, suggesting a convergent evolution of this enzymatic activity [28]. It is thus tempting to speculate that other *BX* genes involved in the metabolism of DIBOA-Glc, such as *BX7, BX13*, and *BX14*, are also derived by independent evolution of wheat and maize.

Major pests of wheat and other species of the grass family (Poaceae) belong to the aphid family (order Hemiptera, family Aphididae), which comprises approximately 5,000 species distributed worldwide [29]. Aphids consume water and nutrients from plants while transmitting toxins through their saliva [30,31], which cause massive yield losses due to both direct and indirect crop damage. They are also responsible for the transmission of 40% of all plant viruses, including the most harmful of plant viruses [32–34]. Other important wheat pests include leaf-chewing insects such as lepidopteran caterpillars. In response to lepidopteran attack, plants massively modify their transcriptome and biosynthesize chemical defense compounds [16,35,36]. While extensive progress has been made on the understanding of the response of grass crops to aphids and caterpillars, little is known about the effects of mite feeding. Recent reports indicated that the two-spotted spider mite (*Tetranychus urticae*) causes damage to cereals such as maize, rice, barley, and wheat, and can be an important pest to these plant species [37,38].

In this study, we aimed to identify *BX6* orthologs in tetraploid and diploid wheat and to study their function in wheat defense. We combined sequence comparisons, phylogenetic analysis, and transcriptome sequencing of tetraploid durum wheat infested with either aphids or caterpillars in order to identify *BX6* genes. Functional gene characterization was performed by applying a virus-induced gene silencing approach to reduce *BX6* expression. *BX6*-silenced plants were then tested in bioassays with different insect herbivores. The results suggest that the BX6-mediated formation of BXDs contributes to plant resistance against insect pests in wheat.

## Materials and Methods

### Plants material and growth conditions

Wheat seedlings were sown in plastic pots, each containing approximately 330 cm^3^ of a tuff mixture with vermiculite (2:1) and an N-P-K fertilizer (20–20–20) [39]. Plants used in this study included hexaploid wheat *Triticum aestivum L*. cultivar Chinese Spring, and tetraploid wheat *Triticum turgidum*, cultivar Svevo. The plants were maintained under controlled growth conditions: 16 h/8 h light/dark; 250–350 μmol photon m^-2^ s^-1^ light intensity from a 3000 lm LED; 24 ± 2°C and watered as needed to maintain moistness [40].

### Phylogenetic analysis

Amino acid sequences of DIBOA-glucoside dioxygenase BX6-like proteins of several grass family species were retrieved from EnsemblPlant website (https://plants.ensembl.org/index.html) and NCBI (https://www.ncbi.nlm.nih.gov/). The multiple sequence alignment was conducted by Clustel-Omega using the default hidden Markov model [41], and the phylogenetic tree was visualized using the MEGA software, version X [42].

### RNA extraction, transcriptome sequencing, and analysis

Tissue samples were collected from the top of the second leaf of *T. turgidum* plants. Samples from two 11-day-old plants were combined into one experimental replicate. Total RNA was extracted using an SV Total RNA Isolation Kit with on-column DNaseI treatment (QIAGEN). Sample integrity was validated using electrophoresis on a 2% agarose gel, and quantified using NanoDrop One-W UV-Vis spectrophotometer (Thermo Fisher Scientific, USA). Then, 2.5 μg of each sample was dried in RNA protective tubes (GenTegra LLC, USA). Library preparation (150 bp read length paired-end) and RNA sequencing was conducted using an Illumina HiSeq 4000 instrument performed by the GeneWIZ Company (www.genewiz.com). Quality control was conducted using FASTQC, and adapters and low-quality sequences were trimmed and removed using Trimmomatic v0.36. Mapping was performed against the *Triticum turgidum* Svevo.v1 reference genome [43] using STAR v2.5.3a [44]. For differential gene analysis, DESeq2 v1.26.0 [45] was utilized to perform differential gene expression analysis (adjusted *p* value < 0.05). TPM values were used for the quantification of gene expression (Supplementary Table S1). The full list of the *IGPS* and *BX* genes in three wheat genotypes, Svevo, Chinses Spring and Zavitan, are presented in Supplementary Table S2. The raw sequence data are available in the NCBI Sequence Read Archive (SRA) as accession PRJNA624178.

### Preparation of mRNA and cDNA for real-time PCR

Leaves were collected, stored at -80 °C, and used for total RNA isolation. Total RNA was isolated using the ZR plant RNA miniprep Kit following the manufacturer’s instructions. DNaseI treatment was given by in column digestion. RNA quantification was done by NanoDrop. For cDNA synthesis, a Verso cDNA synthesis kit was used following the manufacturer’s instructions. For quantitative PCR (qRT-PCR), we quantified the following genes: *TaBX6* (TraesCS2A02G051700, TraesCS2B02G066000, TraesCSU02G024200, TraesCSU02G156500), and glyceraldehyde 3-phosphate dehydrogenase (*GAPC*; gene ID EF592180), which was used as an endogenous control gene. Reactions in a total volume of 10 μl were prepared using Power SYBR Green PCR Master Mix (Thermo Fisher Scientific, MA, USA) following the manufacturer’s instructions, and run on an ABI 7500 Real-Time PCR System (Applied Biosystems, MA, USA) for 40 cycles followed by a dissociation stage, according to standard protocols. The primer list is presented in Supplementary Table S3.

### Transient gene silencing using RNA interference (RNAi)

The transient silencing of genes in hexaploid *T. aestivum*, was carried out by virus-induced gene silencing (VIGS) with the *barley stripe mosaic virus* (BSMV) vector composed of three RNAs (α, β, and γ), which are capped at the 5’ end and form a tRNA-like hairpin secondary structure at the ‘3 terminus [46,47]. A DNA fragment of 230 bp conserved in all four *TaBx6* genes (TraesCS2A02G051700, TraesCS2B02G066000, TraesCSU02G024200, TraesCSU02G156500) was amplified from the cDNA as described above, then cloned into the BSMV system using the restriction enzyme *Apa*I (RNAγ antisense orientation). The primer pair TaBx6_ApaI_BSMV was used for *TaBX6* genes and is shown in Supplementary Table S3. A DNA fragment of *Phytoene desaturase* (PDS) was fused into the BSMV system and employed as a photobleaching phenotype control [47–50]. An empty vector was used as a control. The constructs were transformed into *Agrobacterium tumefaciens* (strain GV3101) and introduced via agroinfiltration into the leaves of the dicot host *Nicotiana benthamiana*, followed by sap extraction from BSMV-infected *N. benthamiana* leaves, which was finally used to rub-inoculate leaves of young Chinese Spring wheat seedlings [47,51–54].

### Benzoxazinoid extraction, analysis, and quantification

Wheat tissue was collected and ground to a fine powder in liquid nitrogen. Then, the frozen powder was weighed and 1:10 (w:v) ratio of extraction solvent was added containing 80% methanol and 0.1% formic acid as previously described [18,19]. In brief, the sample mix was vortexed, the tubes were shaken for 40 min at 4 °C, and centrifuged for 5 min at maximum speed. The supernatant was filtered using a 22 μm filter plate (EMD Millipore Corp., Billerica, MA, USA) for 5 min at 3,000 g and then covered with a WebSeal Mat. Samples were injected into a DIONEX UltiMate 3000 high-performance liquid chromatography (HPLC) system using a C18 reverse-phase Hypersil GOLD column (3 µm pore size, 150×4.60 mm; Thermo Fisher Scientific, Germany). The column oven temperature was set to 40 °C and UV-VIS absorbance spectra at 190-400 nm. The parameters for BXD metabolite separation were applied as previously described [40,55]. For BXD quantification, the chromatograms were compared with the authentic standards and plant crude extract. The two BXDs, DIMBOA and DIBOA, were purchased from Toronto Research Chemicals (Toronto, Canada), and used as authentic standards, as well as two crude extracts, a mix of DIMBOA-Glc:DIM_2_BOA-Glc in a ratio of 80:20, and a mix of HDMBOA-Glc:HDM_2_BOA-Glc in a ratio of 86:14. The calibration curves were calculated by running authentic standards and crude extracts in different concentrations, ranging from 0.5-50 µg/ml. For each BXD compound, peak area was measured using the Chromeleon software and the final concentration was normalized to mg per gram fresh weight. Furthermore, we used ultra-high-performance liquid chromatography – high resolution mass spectrometer (Thermo; Q Exactive mass spectrometer and Ultimate 3000 UPLC) with tandem mass spectrometry (MS/MS) – to fractionate the metabolites into smaller fragment ions and to allow the rebuilding of the molecules’ structure. The samples were run in positive and negative ion modes with mass resolution of 15,000 at 100 to 1000 *m/z*. Chromatographic separation was performed using the same column and mobile phase as mentioned above for HPLC, in a flow rate of 0.8 mL/min. Injection volume was 2µL for both ion modes with the total run time of 18 min. System control and data processing were performed by using the Xcalibur software (Version 4.1), Thermo Fisher, Germany). We identified the following putative BXD metabolites: i) HDMBOA-Glc (2-O-beta-D-glucopyranosyloxy-4,7-dimethoxy- (2H)-1,4-benzoxazin-3(4H)-one and HM2BOA-Glc (2-O-beta-D-glucopyranosyloxy-7,8-dimethoxy-(2H)-1,4-benzoxazin-3(4H)-one) [26,56]; ii) DIMBOA-3 Hex (2,4-dihydroxy-7-methoxy-1,4-benzoxazin-3-one with three hexoses), and iii) HBOA-2 Hex (2-hydroxy-1,4-benzoxazin-3-one with one hexose). Information for each BXD metabolite, including retention time, possible molecular formula, molecular weight, ion fragments in negative and positive ion modes, are presented in Supplementary Table S4.

### Insect treatments and herbivore bioassays

The bird cherry-oat aphids (*Rhopalosiphum padi*), which are phloem-sap feeding insects, were maintained on 2-week-old plants of a *T. aestivum* cultivar named Rotem (Agridera Seeds & Agriculture LTD, Israel) [55]. The colony was reared under the growth regime as described above. The 2^nd^ instar stage Egyptian cotton leafworms (*Spodoptera littoralis*), which are leaf chewing insects, were provided by Dr. Rami Horowitz (ARO). The caterpillars were fed for two days on 2-weeks old *T. aestivum* (cultivar Chinese Spring) seedlings pre-experiment. For the RNAseq analysis, ten adult *R. padi* aphids or three caterpillars were confined to the upper part of the second leaf of *T. turgidum* ssp. *durum* (cultivar Svevo) seedlings using clip cages (4.5 cm in diameter) for 6 h. Control plants held empty cages without insects. For the aphid bioassay, ten adult *R. padi* aphids were confined to the upper part of the second leaf of *T. aestivum* (cultivar Chinese Spring) plants using a clip cage (4.5 cm diameter) and progeny was calculated by counting the total number of aphids (adults and nymphs) after 4 days of infestation (dpi) [40,55]. For the caterpillar performance assay, the second leaves of *T. aestivum* (cultivar Chinese Spring) plants were caged with three larvae (2^nd^ instar) in breathable cellophane bags for 3 days. After this time period, the caterpillar body weight was measured (mg fresh weight). Eggs of the cell-content feeding two-spotted spider mite (TSSM; *Tetranychus urticae*) were obtained from Biobee Sde Eliyahu Ltd and were maintained for least three generations before the experiment, on 4-5 weeks old *T. aestivum* cv. Rotem plants. For the oviposition assay, a TSSM female was placed on the abaxial side of a leaf segment (approximately 4 cm long) and kept on water-saturated cotton wool in a plastic container under growth room conditions. After 3 days, eggs were counted under a binocular microscope [39].

### Statistical analysis

The Student’s *t*-tests were conducted using JMP13 (www.jmp.com, SAS Institute, Cary, NC, USA) and figure presentation was conducted using Microsoft Excel 2010. The RNAseq statistics are described above.

## Results

### Phylogenetic analysis of wheat BX6

A very recent study indicated that hexaploid bread wheat *T. aestivum* (BBAADD) cv Chinese Spring possesses four *BX6* gene paralogs [27]. The sequences of these genes *TaBX6-1-TaBX6-4* are available in GenBank/EMBL/DDBJ databases. A BLAST [57] analysis against the EnsemblPlant database (IWGSC genome assembly) suggested the following loci: *TaBX6-1* (GeneBank - LC519324; EnsemblPlant ID - TraesCSU02G024200), *TaBX6-2* (GeneBank - LC519325; EnsemblPlant ID - TraesCSU02G156500), *TaBX6-3* (GeneBank - LC519326; EnsemblPlant ID - TraesCS2B02G066000), and *TaBX6-4* (GeneBank - LC519327; EnsemblPlant ID - TraesCS2A02G051700). An ortholog of *TaBX6* genes could be identified in the tetraploid (BBAA) wheat *T. turgidum* ssp. *durum* cv. Svevo (EnsemblPlant ID TRITD2Bv1G015490), and also in the genome of wild emmer wheat *T. turgidum* ssp. *dicoccoides* accession Zavitan (TRIDC2BG006360), indicating that *BX6* genes are located on the subgenome B. The *Bx6* orthologs on the A subgenome are not yet annotated. The gene IDs of *BX6* genes in tetraploid and hexaploid wheat are listed in Supplementary Table S2. To identify further *BX6* orthologs/paralogs, the well-characterized BX6 from maize [58] and ScBX6 from rye (*Secale cereale*) [59,60] were used to conduct a TBLASTN search against the genomes of hexaploid, tetraploid, and other Triticeae species, including *T. urartu*, a diploid progenitor of the bread wheat A subgenome, and *Aegilops tauschii*, a diploid progenitor of the bread wheat D subgenome [59]. The progenitor of the B subgenome is yet identified [61], therefore was not included in the analysis. Next, a phylogenetic tree was constructed using an Unweighted Pair Grouping with Arithmetic Mean (UPGMA) clustering method to analyze the evolutionary relationships of the identified potential BX6 proteins in grass species known to accumulate BXDs. As shown in Figure 2, ScBX6 was clustered together with TaBX6-1 and TaBX6-4, from hexaploid wheat, and with BX6 proteins from the two diploid wheat species *T. urartu* (TuBX6) and *Aegilops tauschii* (AtaBX6). While TaBX6-3 formed a cluster together with *T. turgidum* TdBX6 and TtBX6, TaBX6-2 was not part of a cluster, suggesting that it has no orthologs in other wheat species.

**Figure 2.**
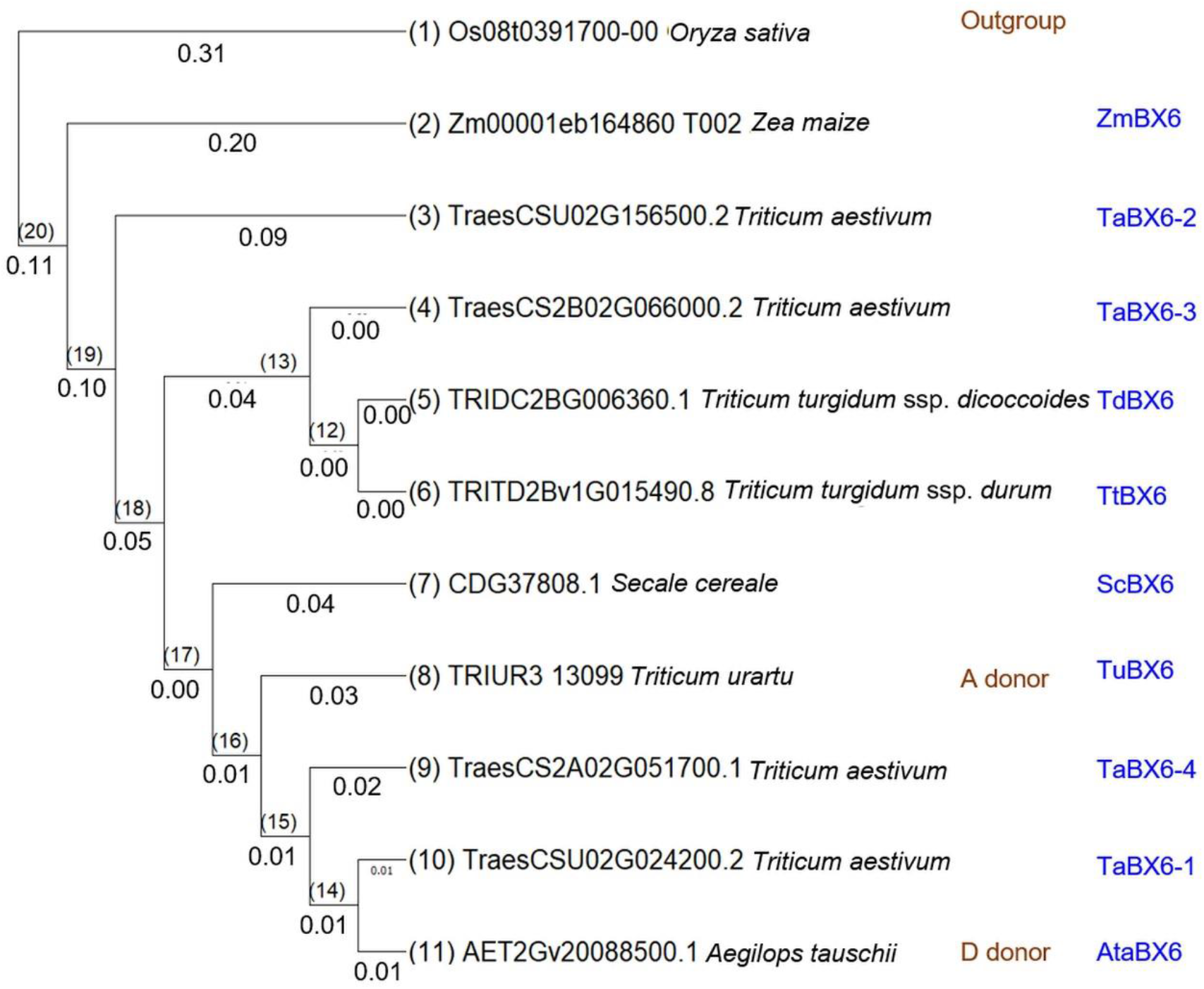
Phylogenetic tree of BX6 protein sequences identified in wheat and other grasses. The phylogenetic tree of BX6 protein sequences includes the closest orthologues to ScBX6 from rye (*Secale cereale*; protein ID: CDG37808.1), maize B73 cultivar (*Zea mays*; protein ID: Zm00001eb164860_T002), *Triticum urartu* (protein ID: TRIUR3_13099), *Aegilops tauschii* (AET2Gv20088500.1), bread wheat Chinese Spring cultivar (*Triticum aestivum;* TraesCS2B02G066000.2, TraesCS2A02G051700.1, TraesCSU02G024200.2, and TraesCSU02G156500.2), durum wheat (*Triticum turgidum* cv. Svevo; TRITD2Bv1G015490.8), and wild emmer wheat Zavitan accession (*T. turgidum* ssp. *dicoccoides*; TRIDC2BG006360.1). As an outgroup, rice from the Japonica Group (*Oryza sativa*; protein ID: Os08t0391700-00) was included. The alignment of protein sequences was performed in Clustal Omega with default parameters, and the tree was generated using the UPGMA method.

### *BX* gene expression is modified in response to insect herbivore feeding

It has been shown that herbivory infestation on maize leaves modifies BXDs metabolism by the upregulation of *BX* gene expression [62,63]. To test whether *BX* genes are also influenced by herbivory in wheat, *T. turgidum* cv. Svevo plants were treated with aphids (*Rhopalosiphum padi*) or caterpillars (*Spodoptera littoralis*) for 6 h using clip cages and transcriptome sequencing was performed. The tetraploid *T. turgidum* was selected because it has only two subgenomes (AABB) and its genome sequence has been recently published [43]. A list of known and putative *BX* genes (see Supplementary Table S2) was created by both searching the literature [7,23,27,28] and the BREADWHEATCYC 2.0 database [64], and by converting the Phytozome12 (https://phytozome.jgi.doe.gov) wheat transcript IDs into IWGSC available on the EnsemblPlant website (https://plants.ensembl.org/index.html). The gene IDs of *BX* orthologs of wild emmer wheat accession Zavitan (WEWSeq_v.1.0) [65], durum wheat (Svevo.v1) [43], and bread wheat (IWGSC) [66] were included. The expression levels of *BX* genes, including the upstream *IGPS* that has recently been shown to catalyze the first committed step of the BXD pathway in maize [13,67] are shown in Table 1. The transcriptome analysis revealed that several *BX* genes were significantly upregulated in response to caterpillar feeding, including *IGPS1, IGPS3, BX2, BX3, BX4, BX8/9*, and *BX10*. In addition, the expression level of *TtBx6* significantly increased and was 4.5 times higher upon caterpillar feeding in comparison to untreated controls. On the other hand, one *BX1* gene (Gene ID: TRITD3Av1G009910) was substantially reduced in response to caterpillar damage, while aphid infestation had no effect on *BX* gene expression (Table 1). We also examined whether the feeding of the abovementioned herbivores affects BXD levels in Chinese Spring leaves. The results in Figure S1 show that both aphids and mites caused no significant changes in BXD levels. Caterpillar attack, on the other hand, induced the levels of several compounds, including HDMBOA-Glc/HM_2_BOA-Glc, DIMBOA-Glc, DIBOA, and HDMBOA-Glc, while DIBOA-Glc and DIMBOA were significantly reduced. These results suggest that the inducibility of BXD levels is dependent on the herbivore species and/or feeding lifestyles.

**Table 1.** List of already characterized and putative *BX* genes in durum wheat. *Triticum turgidum* ssp. *durum* cv. Svevo plants were treated with *Spodoptera littoralis* caterpillars, or *Rhopalosiphum padi* aphids for six hours, followed by gene expression analysis using RNAseq. Gene expression values significantly upregulated upon herbivore feeding relative to untreated control (log2 fold-change [FC] >1) are shown in red. Gene expression values that were significantly downregulated upon herbivore feeding relative to untreated controls (log2 fold-change [FC] <-1) are shown in blue. In bold font, *p* ≤ 0.05, adjusted false discovery rate [FDR]. Grey cells represent gene expression below detection levels.

| Function | Gene short name | Gene ID *Triticum turgidum* Svevo.v1 | log2 FC aphid/con | p adj aphid/con | log2 FC cat/con | P adj cat/con |
| --- | --- | --- | --- | --- | --- | --- |
| Indole-3-glycerol-phosphate synthase | IGPS1 | TRITD5Av1G125990 |  |  | 4.26 | **2.08E-02** |
|  | IGPS1 | TRITD5Bv1G104060 | -0.16 | 1.00E+00 | -0.18 | 8.11E-01 |
|  | IGPS2 | TRITD2Av1G200520 | -0.16 | 1.00E+00 | -0.82 | 4.68E-03 |
|  | IGPS2 | TRITD2Bv1G164620 | -0.16 | 1.00E+00 | -0.34 | 3.70E-01 |
|  | IGPS3 | TRITD7Av1G265410 | 0.33 | 9.81E-01 | 2.74 | **2.88E-11** |
|  | IGPS3 | TRITD7Bv1G217160 | 0.07 | 1.00E+00 | 2.83 | **1.93E-08** |
| Indole-glycerol phosphate aldolase | BX1 | TRITD1Av1G225320 | -0.01 | 1.00E+00 | 0.46 | 1.52E-01 |
|  | BX1 | TRITD1Bv1G223090 | 0.10 | 1.00E+00 | 0.05 | 9.27E-01 |
|  | BX1 | TRITD3Av1G009910 | -0.20 | 9.97E-01 | -1.16 | **3.26E-05** |
|  | BX1 | TRITD3Bv1G010680 | -0.23 | 9.86E-01 | -0.21 | 6.10E-01 |
|  | BX1 | TRITD5Bv1G217360 | -0.20 | 1.00E+00 | -0.28 | 7.23E-01 |
|  | BX1 | TRITD7Av1G211620 | 0.14 | 1.00E+00 | 0.49 | 3.71E-01 |
|  | BX1 | TRITD7Bv1G163280 | -0.24 | 1.00E+00 | -1.06 | 5.15E-02 |
| Indole monooxygenase | BX2 | TRITD4Bv1G126740 |  |  | 6.68 | **5.93E-04** |
|  | BX2 | TRITD3Bv1G020330 | -3.32 | 8.07E-01 | 2.19 | 4.56E-01 |
| Indolin-2-one monooxygenase | BX3 | TRITD2Av1G009970 | -1.85 | 9.07E-01 | 4.94 | **5.42E-07** |
|  | BX3 | TRITD5Av1G003600 | -0.56 | 8.15E-01 | -0.11 | 8.62E-01 |
|  | BX3 | TRITD5Bv1G003050 | -0.42 | 1.00E+00 | -0.14 | 9.04E-01 |
| 3-hydroxyindolin-2-one monooxygenase | BX4 | TRITD2Bv1G006380 | -0.92 | 9.66E-01 | 0.92 | 4.63E-01 |
|  | BX4 | TRITD2Av1G010380 | -0.15 | 1.00E+00 | -0.52 | 1.23E-01 |
|  | BX4 | TRITD0Uv1G013350 | 0.32 | 1.00E+00 | -0.78 | 2.42E-01 |
|  | BX4 | TRITD5Av1G003590 |  |  | 3.34 | 1.72E-01 |
|  | BX4 | TRITD5Bv1G003040 | -0.93 | 8.00E-01 | 0.67 | 4.43E-01 |
|  | BX4 | TRITD5Bv1G170690 | -0.20 | 1.00E+00 | -0.56 | 5.30E-01 |
|  | BX4 | TRITD5Bv1G230110 | 0.29 | 1.00E+00 | 2.86 | **1.99E-02** |
| HBOA monooxygenase | BX5 | TRITD5Bv1G003030 | -1.28 | 9.69E-01 | 3.98 | **1.12E-03** |
| DIBOA 2-D glucosyltransferase | BX8/9 | TRITD6Av1G013160 | -0.55 | 9.95E-01 | 1.10 | 1.65E-01 |
|  | BX8/9 | TRITD6Bv1G019950 |  |  | 5.58 | **3.47E-03** |
|  | BX8/9 | TRITD7Av1G035190 | -0.31 | 1.00E+00 | 0.95 | 1.94E-01 |
|  | BX8/9 | TRITD7Bv1G005150 | -0.26 | 1.00E+00 | 1.08 | **7.58E-03** |
| DIMBOA-glucoside<br>dioxygenase | BX6-3 | TRITD2Bv1G015490 | 0.57 | 9.78E-01 | 2.19 | **2.82E-03** |
| O-methyltransferase | BX10 | TRITD1Bv1G229630 | -0.55 | 1.00E+00 | 3.42 | **4.27E-05** |
|  | BX10 | TRITD1Bv1G229480 | -0.18 | 1.00E+00 | 3.42 | **2.32E-03** |
|  | BX10 | TRITD4Av1G257180 | -1.35 | 4.40E-01 | -0.22 | 8.13E-01 |
|  | BX10 | TRITD2Bv1G031430 |  |  | -2.14 | 3.58E-01 |

### Silencing of *TaBX6* led to reduced BXD accumulation

To investigate the *BX6* gene function *in planta*, we used RNA interference (RNAi), a sequence-specific gene suppression system, to simultaneously silence the *BX6* homologs from all subgenomes (A, B, and D) in bread wheat [48]. A 230 bp of DNA fragment, conserved in all wheat subgenomes, was designed and introduced into the Barley Stripe Mosaic Virus (BSMV) to create a dsRNA-producing construct (*BSMV::bx6*). The *T. aestivum* cv. Chinese Spring was infected with this construct or *BSMV::empty* (control). Plants infected with *BSMV::bx6* showed reduced *TaBX6* transcript levels in comparison to *BSMV::empty* plants, as presented in Figure 3A. Moreover, methanol extraction of leaf tissue and subsequent liquid chromatography showed that downregulation of *TaBX6* gene expression resulted in a significantly reduced accumulation of DIM_2_BOA-Glc, DIMBOA-Glc, and HDMBOA-Glc (Figure 3B). Two putative BXDs, DIMBOA-3-Hex and HBOA-2-Hex, were also lower in *BMSV::bx6* relative to *BSMV::empty* leaves, while the HDMBOA-Glc/HM_2_BOA-Glc levels remained unchanged. Overall, our results demonstrated that a reduction in *TaBX6* transcript levels affects BXD biosynthesis in wheat.

**Figure 3.**
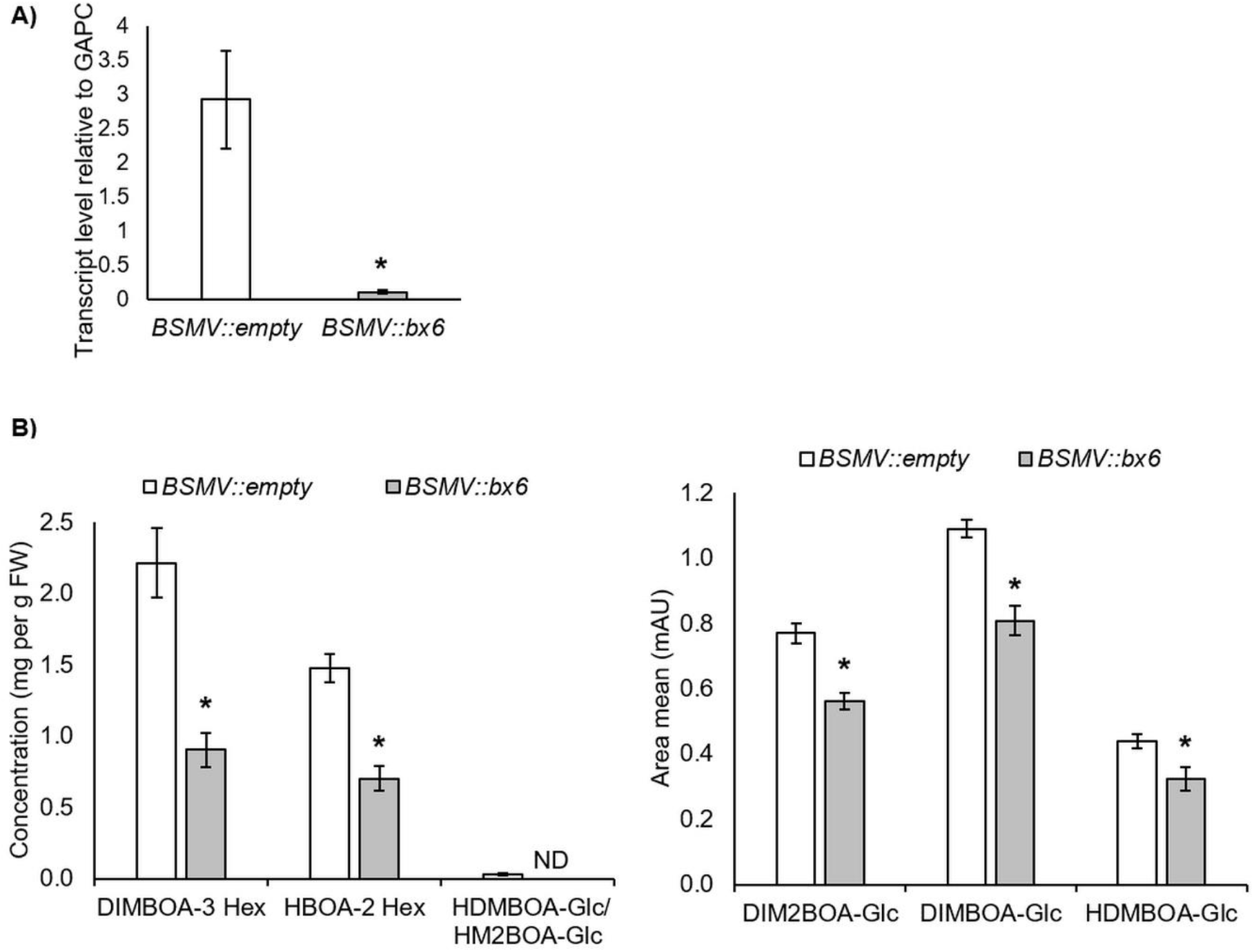
Effect of *TaBX6* gene silencing on benzoxazinoid levels in wheat. Five-days old *T. aestivum* cv. Chinese Spring plants were infected with *BMSV::empty* (grey bars), or *BMSV::bx6* (grey bars), VIGS constructs. After 14 days, leaves were collected for analysis. A) Gene expression of *TaBx6* relative to the reference gene glyceraldehyde 3-phosphate dehydrogenase (*GAPDH*) was quantified using qRT-PCR (n = 11-12). B) BXD accumulation was analyzed by HUPLC. Known BXDs are shown in mg per g fresh weight (middle panel) and putative BXDs are shown in peak area mean (mAU) (left panel). Asterisks indicate significant differences, \**p* < 0.05, Student’s *t*-test (n = 5-7).

### *TaBX6* contributes to plant defense against insect herbivores

To understand the biological role of *TaBX6* in plant defense against insects, we tested whether the reduction of *TaBX6* expression and BXD accumulation affect pest fitness parameters. In a first bioassay, we studied the response of the aphid *R. padi* to a reduction in BXDs in Chinese Spring seedlings infected with *BMSV::bx6* compared to *BMSV::empty*. Figure 4A shows that the total numbers of nymphs and adult aphids were significantly increased in the *BSMV::bx6* plants relative to the empty vector controls. In the second bioassay, we examined the weight gain of *Spodoptera littoralis* caterpillars grown on *BMSV::bx6* and *BMSV::empty* plants. Three caterpillars were applied to *BX6*-silenced and control plants and their body weight was measured after three days. The results presented in Figure 4B reveal a slight increase in caterpillar weight after feeding on *BSMV::bx6* plants. However, this induction in body weight was not statistically significant compared to the empty vector plants. The third biological test was conducted with the two-spotted spider mite *T. urticae*. We applied one adult female on the abaxial page of leaf segments infected with VIGS, and counted the number of eggs laid by the mite. This bioassay found that the average number of eggs laid by the mites was significantly higher on the *BSMV::bx6* plants than the empty vector-infected plants (Figure 4C). Altogether, our findings showed that low *TaBX6* expression levels (Figure 3A) led to a reduction in BXD accumulation (Figure 3B) and consequently increased pest susceptibility (Figure 4).

**Figure 4.**
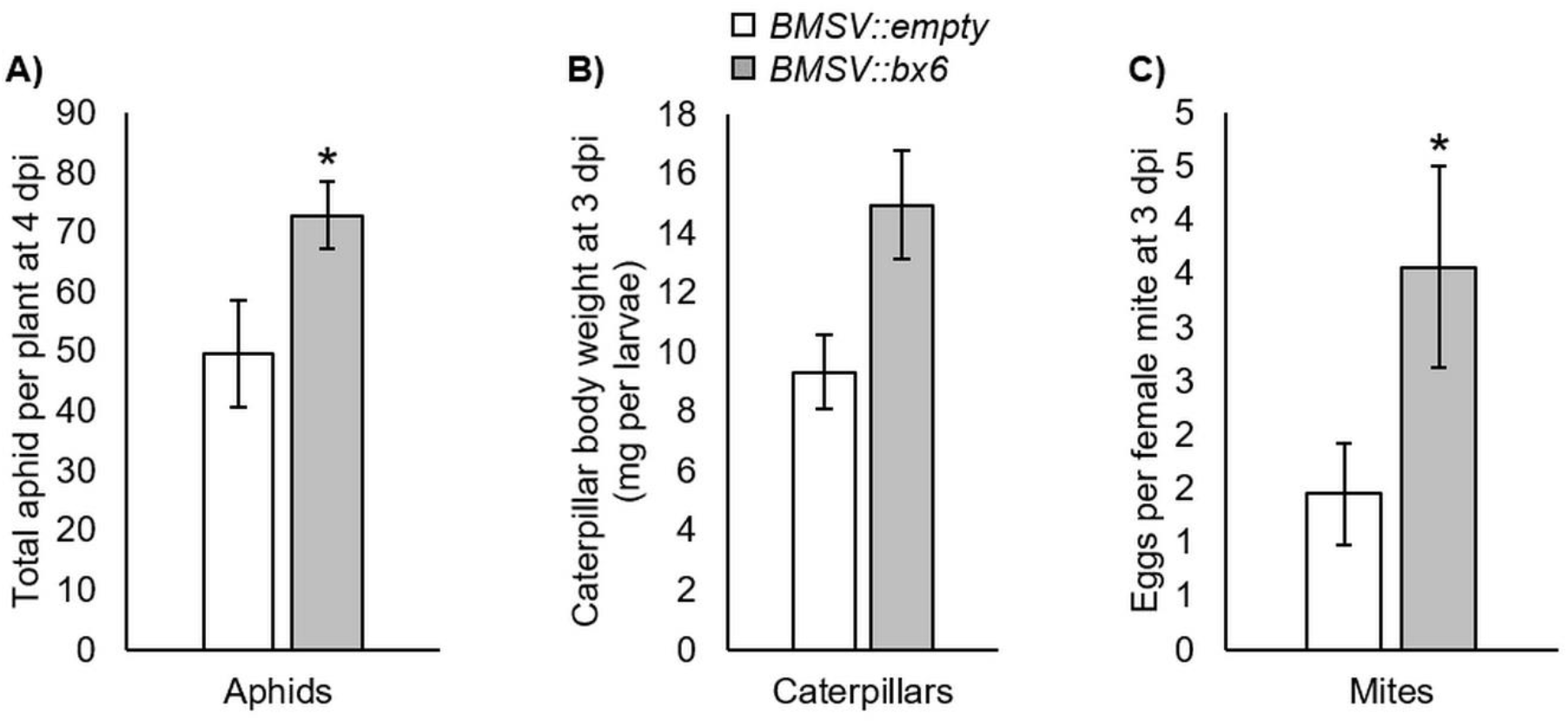
Bioassays with different insect herbivores and *BX6* silenced *T. aestivum* plants. Five day-old Chinese Spring wheat seedlings were infected with *BMSV::bx6* (white bares), or with *BMSV::empty* (grey bars), for six days and then exposed to different insect pests. A) Ten *Rhopalosiphum padi* aphids were applied on the plants and the mean number of aphids per plant was counted after four days from infection (n=10). B) *Spodeptera littoralis* caterpillars were applied to plants and after 3 dpi the larve’s body weight was measured (n= 9-10). C) The oviposition rate of one adult *Tetranychus urticae* female mite per plant was determined on leaf segments. The figure shows the average total mite egg production after 3 dpi (n=12). Asterisks indicate significant differences between *BMSV::bx6* and *BMSV::empty* plants as control (Student’s *t*-test; \**p* < 0.05; mean ± SE).

## Discussion

Benzoxazinoids are specialized metabolites that play diverse roles in plant defense against herbivores and pathogens. Their formation and function have been extensively studied in maize (Figure 1), and recently in wheat and rye [12,15,68]. Wheat is the staple crop for an estimated 35% of the world’s population. During 2016-2018, 68% of global production was used for human consumption, 20% for livestock feed, and 12% for biofuel and other purposes [69]. Therefore, elucidating the BXD biosynthesis pathway in wheat is critical for improving the natural mode of defense against herbivores and to ensure food security. In the present study, TtBX6 from cultivated tetraploid wheat and TdBX6 from the wild emmer wheat were identified using sequence similarity to already known BX6 orthologs in maize (*Zea mays*) [58], rye (*Secale cereale*), and *T. aestivum* [27,59,60]. Sequence comparisons showed that *S. cereale* ScBX6 has a greater similarity with wheat BX6 paralogs [60] than with maize BX6 (Figure 2). In hexaploid wheat, four *TaBX6* genes were identified. *TaBX6-4* was found to be located on subgenome A, while *TaBX6-3* is located on subgenome B. The genomic locations of the two remaining genes *TaBX6-1* and *TaBX6-2* (are still unknown. The high sequence similarity of about 98% between TaBX6-1 and *A. tauschii* AtaBX6 might indicate that *TaBX6-1* is the ortholog of *AtaBX6* and is located on subgenome D. Transcriptome sequencing of durum wheat infested with *S. littoralis* caterpillars for 6 h showed that 13 *BX* and *IPGS* genes were upregulated, including *TtBX6* (Table 1). However, aphid feeding had no effect on *BX* gene expression. Very recent research published by Sue *et al*. (2021) reported on the identification and biochemical characterization of four *TaBX6* genes in hexaploid wheat; however, the function was not described *in planta*. In the present study, a virus-induced gene silencing (VIGS) approach was used, which frequently aids in understanding gene function under biotic and abiotic stresses [70,71]. In wheat, for example, this approach has been used to target a wide range of gene functions, including starch, fatty acid, and α-gliadin biosynthesis [72–74]. As a result of *TaBX6* gene silencing, the amounts of DIM_2_BOA-Glc, DIMBOA-Glc, and HDMBOA-Glc were reduced and the plants were more susceptible to aphids and mites (Figures 3-4). Overall, this suggests that the dioxygenase TaBX6 is involved in the defense responses of wheat seedlings against herbivory.

The intensity of plant defense responses differ by herbivore lifestyles and are essential for maintaining plant fitness [28]. BXDs are involved in plant defense against caterpillars (leaf chewing insects). For example, exposure of *S. littoralis* and *S. frugiperda* caterpillars on maize seedlings strongly increased DIMBOA-Glc methylation and the production of HDM_2_BOA-Glc [36]. BXDs also contribute to defense against aphids (phloem-sap feeding insect) by possessing antifeedant and antibiosis properties. For example, DIMBOA is associated with callose formation [9], whereas HDMBOA-Glc was shown to have toxic effects [17]. In wheat and maize leaves, constitutive levels of BXDs are involved in determining aphid performance, while inducible changes were minor [17,26]. The results in this study revealed massive changes in *BX* and *IGPS* gene expression levels upon caterpillar feeding for 6 h, while there were no significant changes after aphid feeding (Table 1). These findings were supported by previous transcriptomic analyses of *BX* genes in maize leaves, which showed that these genes were dramatically upregulated upon *S. exigua* caterpillar attack [63], but only mildly upregulated upon corn leaf aphid (*Rhopalosiphum maidis*) infestation [75]. Sue *et al*. (2021) treated hexaploid wheat seedlings with jasmonic acid or grew them in dark conditions (etiolation) and found only induction of DIMBOA-Glc, while none of the *BX* genes (*BX1-BX6* and *BX8*) were upregulated. This suggests that the correlation between *BX* gene expression levels and BXD accumulation differ between stresses and that other factors might determine the levels of BXDs in wheat.

Plants with mutated *BX* genes have been used to discover the role of benzoxazinoids in plant defense. For example, *BX1, BX2*, and *BX6 Ds* transposon insertion mutants in maize inbred line W22 have shown a reduction in BXD levels and enhanced susceptibility to *R. maidis* aphids and increased two-spotted spider mite progeny, where these effects were stronger in *bx1::Ds* and *bx2::Ds* mutants than *bx6::Ds* [75,76]. This suggests a substantial functional redundancy at the respective step in the BXD pathway; for instance, the partial toxicity of DIBOA-Glc that is still produced in the *bx6* mutant [76]. In our research, the *TaBX6* gene silencing resulted in a significant induction of both aphid reproduction and two-spotted spider mite oviposition relative to controls, while the caterpillar body weight was only slightly (and not significantly) increased (Figure 4). The impact of BXD deficient maize mutants (*bx1::Ds* and *bx2::Ds*) on pest feeding suggests stronger susceptible phenotypes of aphids and mites than caterpillars. Aphid reproduction on *bx1::Ds* and *bx2::Ds* mutants was approximately 7 times higher than wildtype [75], mite progeny on *bx1::Ds* and *bx2*::*Ds* mutants were approximately 3.5 and 6 time higher than wildtype [76], respectively, and caterpillar body weights on *bx1::Ds* and *bx2::Ds* mutants were approximately 1.4 higher than wildtype [63]. Here we suggest that in Chinese Spring wheat seedlings, the BXDs are less effective against caterpillar feeding than aphids and mites. Altogether, the biosynthesis, regulation, and function of these compounds on herbivores are species-specific.

The two tetraploid genotypes, durum wheat (*T. turgidum* ssp. *durum*) and wild emmer (*T. turgidum* ssp. *dicoccoides*) have been sequenced [43,65], which allows us to use them for gene discovery. Comparing *TaBX6*(1-4) to these tetraploid wheat genotypes indicated that *BX6* genes are located on subgenome B, while the homologs on subgenome A are known (Table S3). The phylogenetic tree indicated that *TaBX6-3* is highly similar to *TtBX6* and *TdBX6*, as all are located on subgenome B (Figure 2). Moreover, the transcriptome analysis of *BX* and *IGPS* genes in Svevo leaves revealed that 13 genes out of 35 were significantly affected by caterpillar feeding; four of them are located on subgenome A and nine located on subgenome B (Table 1). This is supported by a previous study that compared the transcript levels of the three hexaploid wheat genomes, which showed that the *BX* homologs located on the B subgenome are the main contributors to the BXD biosynthesis, especially in shoots [23,77]. Thus, we suggest that *BX6* located on subgenome B is predominantly involved in the BXD biosynthesis.

## Conclusion

Insect herbivory often causes severe damage to plants, resulting in significant losses in crop yield. The gradual increase in global temperatures has promoted the expansion of pest populations to new regions and increased their reproduction rate [78]. Therefore, it is of the utmost importance to rigorously explore natural plant defense mechanisms and traits to discover additional modes of resistance that could be deployed against pests. In this study, we identified the *BX6* gene and its function in BXD biosynthesis in diploid and tetraploid wheat. We used the VIGS approach, which is used frequently to dissect gene function under biotic and abiotic stresses. However, its effectiveness is limited to a short time window after infection. To further elucidate the function of *BX6*, we suggest generating stable mutants and testing their effect on herbivory in combination with other biotic and abiotic stresses.

## Acknowledgments

We are grateful to Kostya Kanyuka (Rothamsted Research, UK) for providing the BMSV cloning vector collection; Zahevit Dadon-Yegana and Noga Sikron Peres (BGU) for their assistance with the HPLC and LC-MS analysis; Nati Weinblum (BGU) for helping with the TSSM mite bioassay; Biobee Sde Eliyahu Ltd. for providing the TSSM mites and Rami Horowitz from the Agriculture Research Organization for providing the caterpillars. We also want to thank Matthias Erb (University of Bern, Switzerland) for kindly providing the benzoxazinoid crude extract and Assaf Distelfeld (University of Haifa) for providing the seeds.

## Supporting Information

**Table S1**. Total RNA-seq values after rlog normalization. Genes were annotated to the A and B subgenomes or unidentified subgenome (U). Differentially expressed genes (DEGs) were obtained using the DESeq2 from R package. Fold changes are presented in log2, *p* value, and adjusted *p* value.

**Table S2**. The full list of *IGPS* and *BX* genes in three wheat genotypes: Svevo, Chinese Spring, and Zavitan. The data includes genes from subgenomes A, B, D, and U (not classified).

**Table S3**. Full list of the primers that were used in this study. Underline represents the cutting area of restriction enzymes.

**Table S4**. Retention time, molecular weight, fragments, and formula of the BXD that were identified using tandem mass spectrometry.

**Figure S1**. BXD induction by insect feeding in Chinese Spring wheat leaves. A) Aphid infestation for 4 d. B) Caterpillar feeding for 3d. C) Mite infestation timepoint bioassays. Asterisks indicate a significant difference (Dunnett test, *p* < 0.05 by (n=5-8)).

