## Supplementary figures and images for "The wheat dioxygenase BX6 is involved in the formation of benzoxazinoids *in planta* and contributes to plant defense against insect herbivores"

### Figure S1

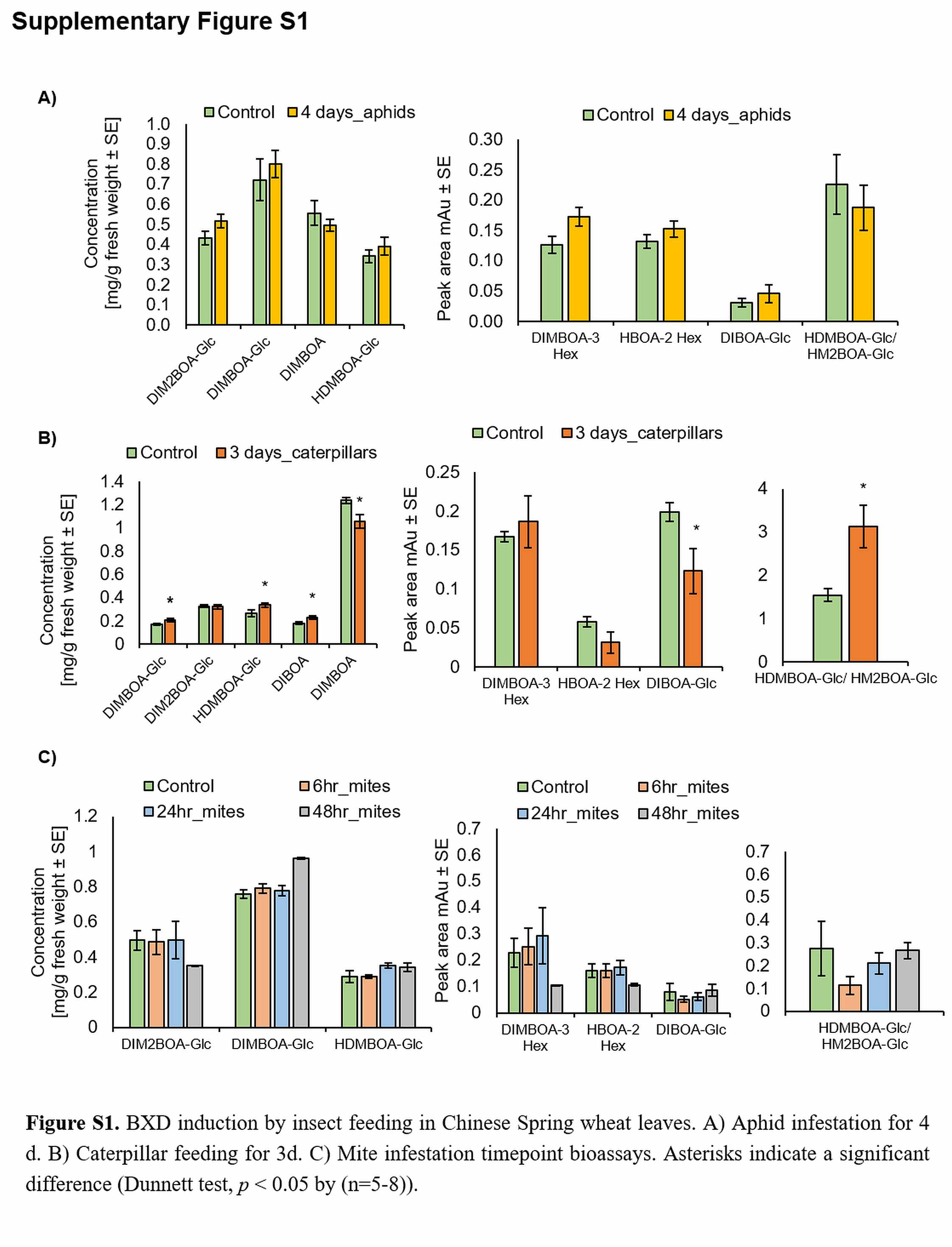
